# The protective association of endogenous immunoglobulins against sepsis mortality is restricted to patients with moderate organ failure

**DOI:** 10.1101/116665

**Authors:** Ignacio Martin-Loeches, Arturo Muriel-Bombín, Ricard Ferrer, Antonio Artigas, Jordi Sole-Violan, Leonardo Lorente, David Andaluz-Ojeda, Adriele Prina-Mello, Ruben Herrán-Monge, Borja Suberviola, Ana Rodriguez-Fernandez, Pedro Merino, Ana M Loza, Pablo Garcia-Olivares, Eduardo Anton, Eduardo Tamayo, Wysali Trapiello, Jesús Blanco, Jesús F Bermejo-Martin, GRECIA Group

## Abstract

**Background:** pre-evaluation of endogenous immunoglobulin levels is a potential strategy to improve the results of intravenous immunoglobulins in sepsis, but more work has to be done to identify those patients who could benefit the most from this treatment. The objective of this study was to evaluate the impact of endogenous immunoglobulins on the mortality risk in sepsis depending on disease severity.

**Methods:** this was a retrospective observational study including 278 patients admitted to the ICU with sepsis fulfilling the SEPSIS-3 criteria, coming from the Spanish GRECIA and ABISS-EDUSEPSIS cohorts. Patients were distributed into two groups depending on their Sequential Organ Failure Assessment score al ICU admission (SOFA < 8, n = 122 and SOFA ≥ 8, n = 156) and the association between immunoglobulin levels at ICU admission with mortality was studied in each group by Kaplan Meier and multivariate logistic regression analysis.

**Results:** ICU / hospital mortality in the SOFA < 8 group was 14.8% / 23.0%, compared to 30.1 % / 35.3% in the SOFA ≥ 8 group. In the group with SOFA < 8, the simultaneous presence of total IgG <407 mg/dl, IgM < 43 mg/dl and IgA < 219 mg/dl was associated to a reduction in the survival mean time of 6.6 days in the first 28 days, and was a robust predictor of mortality risk either during the acute and the post-acute phase of the disease (OR for ICU mortality: 13.79; OR for hospital mortality: 7.98). This predictive ability remained in the absence of prior immunosupression (OR for ICU mortality: 17.53; OR for hospital mortality: 5.63). Total IgG <407 mg/dl or IgG1 < 332 mg/dl was also an independent predictor of ICU mortality in this group. In contrast, in the SOFA ≥ 8 group, we found no immunoglobulin thresholds associated to neither ICU nor to hospital mortality.

**Conclusions:** endogenous immunoglobulin levels may have a different impact on the mortality risk of sepsis patients based on their severity. In patients with moderate organ failure, the simultaneous presence of low levels of IgG, IgA and IgM was a consistent predictor of both acute and post-acute mortality.

## Background

Pre-evaluation of endogenous immunoglobulin levels has been proposed as a potential tool to identify sepsis patients deserving replacement treatment with intravenous immunoglobulins (IVIG), to improve its results in this severe condition [1]. Nonetheless, the impact of endogenous immunoglobulins levels on the risk of mortality in sepsis remains a controversial issue. A recent meta-analysis leaded by Shankar-Hari M found that the prevalence of IgG hypogamma-globulinaemia on the day of sepsis diagnosis is as high as 70%, but this finding did not identify a subgroup of patients with a higher risk of death [2]. Recently, results from the SBITs (Score-based immunoglobulin G therapy of patients with sepsis) study, showed that initial low IgG levels did not discriminate between survival and non-survival in patients with severe sepsis and septic shock [3]. In addition, patients with the highest IgG levels (fourth quartile) showed a statistically significant higher mortality in a risk-adjusted calculation compared to the reference quartile [3]. A previous report from our group supported that the answer could be in considering immunoglobulin isotypes not as isolated entities but in evaluating their prognostic ability in combination [4] [5].

There are a number of factors that in our opinion have not been appropriately addressed in the studies evaluating the predictive ability of immunoglobulins: 1) we have demonstrated in a recent article that disease severity strongly influences biomarker performance in sepsis [6]; 2) the influence of previous immunosupression has not been evaluated [5]; 3) the impact of immunoglobulins on hospital mortality has not been sufficiently studied, with the majority of works being focused on the acute period of the disease [2]; 4) finally, the ability of endogenous immunoglobulin levels to predict mortality in patients fulfilling the SEPSIS-3 criteria [7] [8] has not been reported to the present moment.

Aimed by the need of identifying patient subsets that could benefit the most from IVIG therapy, we have now evaluated the ability of endogenous immunoglobulins levels and also of a combined immunoglobulin score to predict mortality risk of sepsis patients fulfilling the SEPSIS-3 diagnostic criteria in two different scenarios of disease severity, at the short and the long term, with and without presence of previous immunosupression.

## Methods

### Study design

patients from two multi-center epidemiological studies on sepsis were merged to evaluate, in a retrospective manner, the association between levels of endogenous immunoglobulins in plasma and mortality depending on disease severity at ICU admission. 180 patients came from the GRECIA study [9], (Grupo de Estudios y Análisis en Cuidados Intensivos) and 98 came from the ABISS-Edusepsis study (AntiBiotic Intervention in Severe Sepsis) [10]. In both studies, patients had sepsis at the time of admission to the ICU. For this study, only those patients fulfilling the new definition proposed by the SEPSIS-3 Consensus were considered [7]. Patients with human immunodeficiency virus (HIV) infection and those undergoing radiotherapy or receiving immunosuppressive drugs, including chemotherapy or systemic steroids, in the last 3 months prior to admission to the ICU were considered to be immuno-suppressed. Exclusion criteria were cardiac arrest, therapeutic effort limitation and lack of informed consent. A common data sheet was developed to collect the clinical data, including medical history, physical examination and haematological, biochemical, radiological and microbiological investigations from the two studies. Treatment decisions were not standardized for all patients but were made by the treating physician, always based upon the Surviving Sepsis Campaign guidelines recommendations.

### Immunoglobulin quantification

a 5-mL sample of blood was collected in an EDTA tube from all patients in the first 12 hours following ICU admission. The blood was centrifuged, and plasma was obtained and stored at − 80 °C until required for immunoglobulin quantification. Plasma levels of immunoglobulins were measured using a multiplex immunoglobulin isotyping kit (Biorad TM, Hercules, CA, USA) on a Luminex platform. All plasma samples from the two studies (GRECIA and ABISS) were tested for immunoglobulin concentrations using the same equipment, to avoid potential bias due to multiplatform testing.

### Statistical analysis

for the demographic and clinical characteristics of the patients, differences between groups were assessed using the χ2 test for categorical variables and the Mann-Whitney U test for continuous variables when appropriate. Patients were split into two groups based upon the percentile 50 for the Sequential Organ Failure Assessment (SOFA) score at ICU admission. In consequence, two groups of patients were generated (SOFA < 8 and SOFA ≥ 8). Deciles of immunoglobulin concentrations were used to categorize patients below or above each decile, creating the corresponding categorical variables. Deciles were calculated for the entire cohort, since no differences for immunoglobulin levels were found between patients with SOFA score <8 and those with SOFA scores ≥ 8 (Table 1). We determined the occurrence of death in each severity group using Kaplan-Meier curves. Time was censored at day 28 following admission to the ICU for this analysis. The first decile showing significant differences between groups based on the log-rank test was considered as the immunoglobulin threshold. We established immuno-scores (ISC) for identifying those patients with the combined presence of low levels of two or more immunoglobulins (below each respective threshold). An overall score of 0 was assigned to all patients with levels below the thresholds for all immunoglobulins forming each immuno-score and a score of 1 to the remaining patients. The dichotomous variables created for each immunoglobulin using the identified thresholds as well as the immuno-scores were further introduced into a multivariate logistic regression analysis to determine the association between immunoglobulin levels and the risk of mortality at the ICU and also at the hospital. Those variables of Table 1 yielding *p* values < 0.1 in the univariate analysis were considered as potential confounding factors and were further introduced in the multivariate one as adjusting variables. Data analysis was performed using SPSS for WINDOWS version 22.0 software (IBM-SPSS, Chicago, IL, USA).

## Results

### I. Clinical characteristics of the patients depending on disease severity at ICU admission

in order to evaluate the potential differences between the patients included in the two severity groups, a descriptive table was built (Table 1). This table is important to find out whether or not patients’ characteristics could explain the different results found for the two groups regarding the association between immunoglobulin levels and the risk of mortality, as showed later in this section. Table 1 shows that patients were elderly individuals, with the group of patients with SOFA score ≥ 8 having a significant higher proportion of men. The most frequent co-morbidities found in both groups of patients were chronic cardiovascular disease, chronic respiratory disease, chronic renal failure and diabetes mellitus. The most severe group of patients showed a significant higher frequency of patients with history of chronic hepatic failure. The proportion of patients with prior immunosupression did not differ in a significant manner between both groups. Sources of infection were similar in the two groups compared, with predominance of sepsis of respiratory, abdominal and urological origin. Both groups presented also a similar proportion of infections caused by Gram + bacteria and fungi, but the most severe group had a significant higher frequency of infections caused by Gram - bacteria along with an overall higher frequency of patients with microbiologically confirmed infection. 94 % of the patients with SOFA ≥ 8 presented with cardiovascular dysfunction, compared with 63% in the group of patients with SOFA < 8. As expected, patients in the most severe group showed higher APACHE-II scores, higher creatinine levels in plasma, and lower platelets counts in blood. In addition, mortality at the ICU and also at the hospital was markedly higher in this group. Immunoglobulin and albumin levels in plasma did not differ in a significant manner between both groups of patients. In those patients with positive microbiological identification, proportion of patients receiving appropriated antibiotic treatment based on the antibiogram results was similar between both severity groups (80% *vs* 73%, *p* = 0.420).

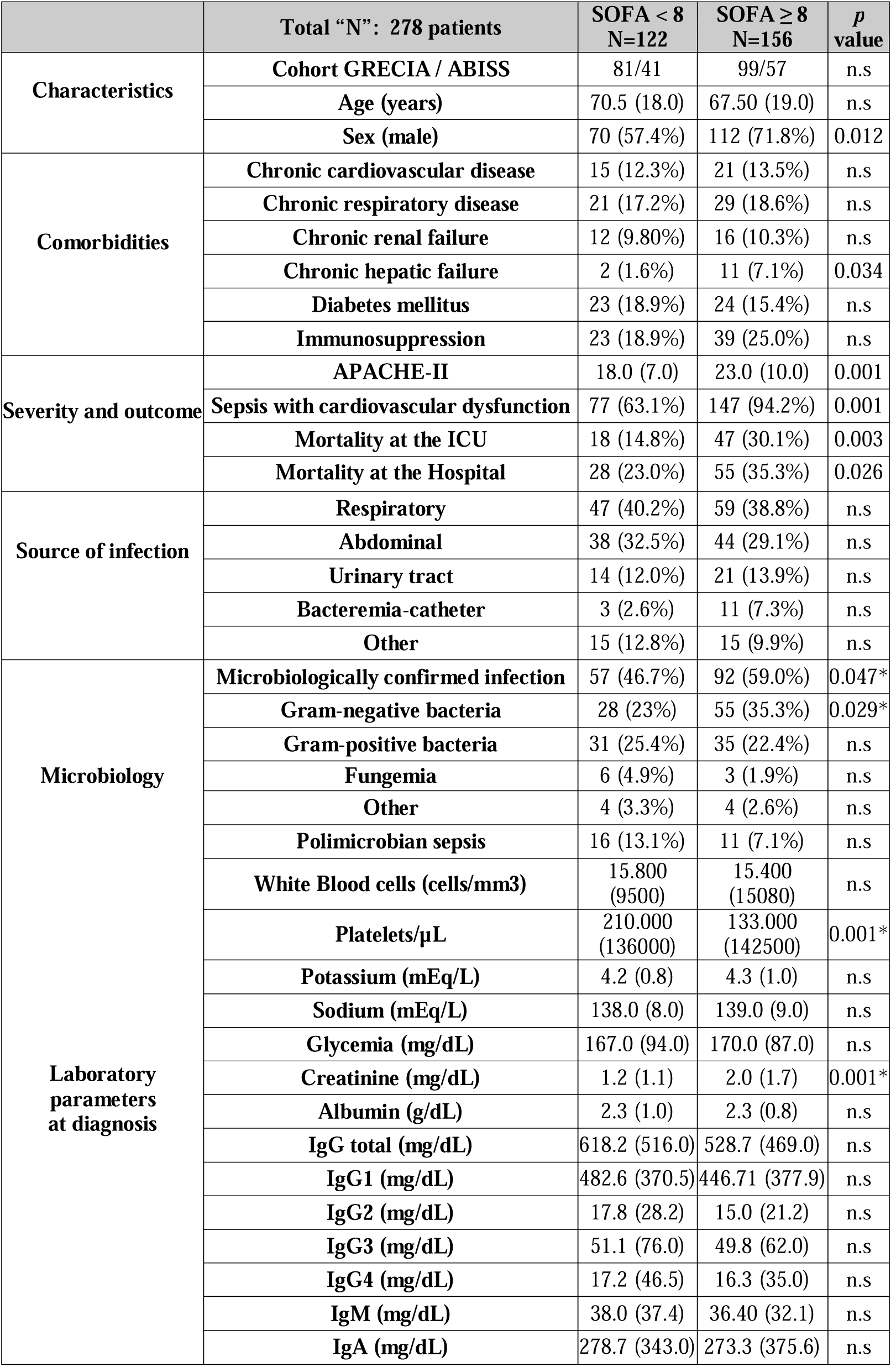
Clinical characteristics of the patients. Continuous variables are shown as median (inter-quartile rank) and categorical variables as n (%). N.s: not significant.

### II. Identification of immunoglobulin thresholds associated with mortality

Kaplan-Meier analysis identified five immunoglobulin thresholds associated with mortality in the group of patients with SOFA < 8, corresponding to IgG, IgG1, IgG2, IgM and IgA (Figure 1). The triple ISC built based upon the thresholds corresponding to the three major immunoglobulin isotypes (IgG, IgM and IgA) showed the highest impact on survival mean time, which was reduced 6.6 days in average in those patients showing a ISC IgGAM = 0 (Figure 1 and Additional file 1). When Kaplan Meier analysis was repeated for those patients of the most severe group (those with SOFA ≥ 8), no thresholds associated with mortality were identified for individual immunoglobulins (Figure 2). In addition, the ISC IgGAM failed to show any association with mortality in this analysis (Figure 2).

**Figure 1:**
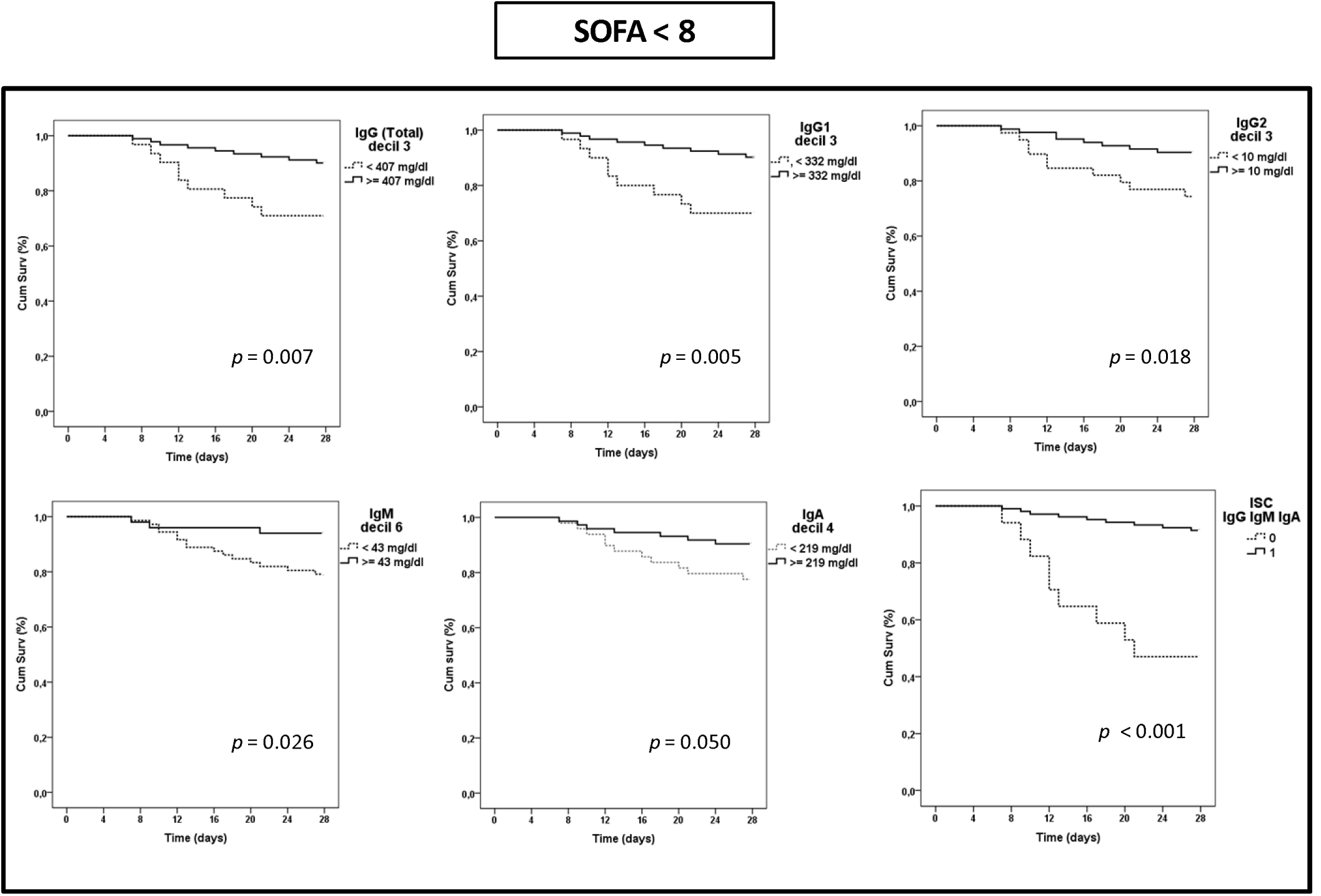
Kaplan Meier analysis for survival depending on immunoglobulin levels in the group with SOFA < 8.

**Figure 2:**
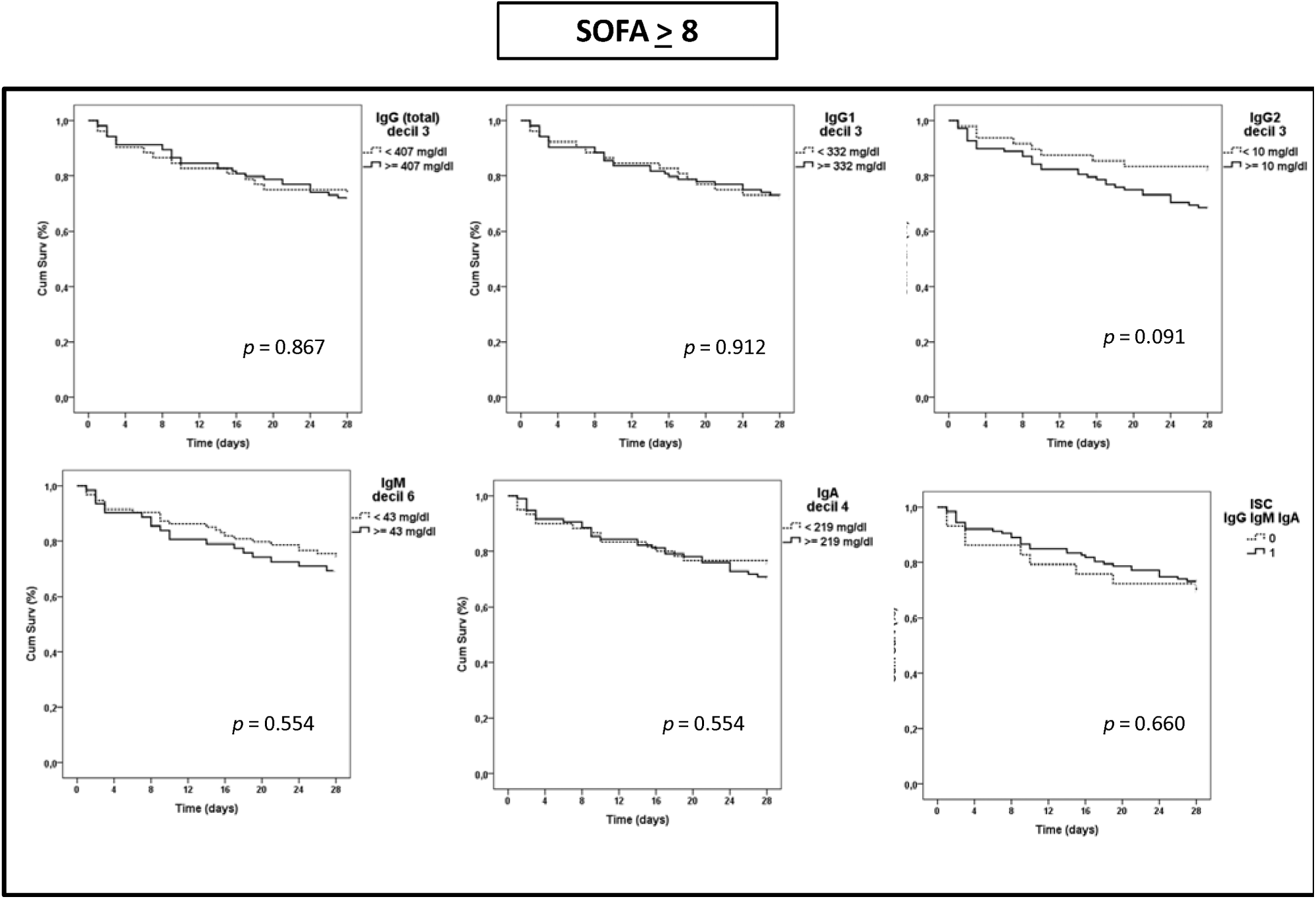
Kaplan Meier analysis for survival depending on immunoglobulin levels in the group with SOFA ≥ 8.

### III. Regression analysis for prediction of mortality risk

to evaluate the impact of the immunoglobulin thresholds on the mortality risk over the acute phase of the disease and also over the post-acute phase of the disease, regression analysis for predicting either ICU and hospital mortality were performed in the group of patients with SOFA < 8:

#### III.1 ICU mortality

we first compared the proportion of survivors and non survivors at the ICU in the patients with immunoglobulin levels below/above these thresholds (Table 2):

**Table 2:** Proportion of survivors and non survivors at the ICU in the group of patients with SOFA score < 8 depending on the immunoglobulin thresholds.

|  |  | *SOFA < 8: ICU mortality* |  |  |  |  |  |
| --- | --- | --- | --- | --- | --- | --- | --- |
|  |  | *Patients with prior immunosuppression included (n=122)* |  |  | *Patients with prior immunosuppression excluded (n=99)* |  |  |
|  | Thresholds | Non survivors / Total "n" in each category | % | *p* | Non survivors / Total "n" in each category | % | *p* |
| **IgG (total)** | < 407 mg/dl | 8 / 31 | 25.8% | 0.045 | 6 / 24 | 25.0% | 0.048 |
|  | ≥ 407 mg/dl | 10 / 91 | 14.8% |  | 7 / 75 | 9.3% |  |
| **IgG1** | < 332 mg/dl | 8 / 30 | 26.7% | 0.034 | 6 / 24 | 25.0% | 0.048 |
|  | ≥ 332 mg/dl | 10 / 92 | 14.8% |  | 7 / 75 | 9.30% |  |
| **IgG2** | < 10 mg/dl | 9 / 39 | 23.1% | 0.076 | 8 / 32 | 25.0% | 0.016 |
|  | ≥ 10 mg/dl | 9 / 83 | 10.8% |  | 5 / 67 | 7.5% |  |
| **IgM** | < 43 mg/dl | 13 / 72 | 18.1% | 0.217 | 9 / 54 | 16.7% | 0.254 |
|  | ≥ 43 mg/dl | 5 / 50 | 10.0% |  | 4 / 45 | 8.9% |  |
| **IgA** | < 219 mg/dl | 10 / 49 | 20.4% | 0.149 | 7 / 39 | 17.9% | 0.253 |
|  | ≥ 219 mg/dl | 8 / 73 | 11.0% |  | 6 / 60 | 10.0% |  |
| **ISC IgGAM** | = 0 | 7 / 17 | 41.2% | 0.001 | 5 / 12 | 41.7% | 0.002 |
|  | = 1 | 11 / 105 | 10.5% |  | 8 / 87 | 9.2% |  |

Those thresholds showing differences at the level *p* < 0.1 in the Table 2 were further evaluated for their association with the risk of ICU mortality by using a multivariate analysis (Table 3). This analysis evidenced that exhibiting levels of IgG, IgG1 below their respective thresholds or having an ISC IgGAM = 0 was a robust, independent risk marker of ICU mortality. The highest Odds Ratio corresponded to the ISC IgGAM. While the presence of patients with prior immunosupression did not affect the predictive ability of IgG, IgG1 and ISC IgGAM, it did influence IgG2, which was only able to predict mortality in those patients with no previous antecedents of immunosupression.

**Table 3.** Multivariate logistic regression analysis to evaluate the association between immunoglobulins and the risk of mortality at the ICU in the group of patients with SOFA score < 8. Adjusting variables for *A.* were [diabetes mellitus] [presence of prior immunosupression] [APACHE-II score] [presence of respiratory infection], [microbiologically confirmed infection]. Adjusting variables for *B.* were [APACHE-II score] [presence of respiratory infection], [presence of abdominal infection] [microbiologically confirmed infection].

|  | SOFA < 8 |  |  |  |  |  |  |  |
| --- | --- | --- | --- | --- | --- | --- | --- | --- |
|  | *A. Patients with prior immunosuppression included (n=122)* |  |  |  | *B. Patients with prior immunosuppression excluded (n=99)* |  |  |  |
| Immunoglobulins | OR | CI 95% |  | *p* | OR | CI 95% |  | *p* |
| IgG (total) < 407 mg/dl | 7.29 | 1.62 | 32.79 | 0.010 | 9.02 | 1.45 | 56.30 | 0.019 |
| IgG1 < 332 mg/dl | 8.02 | 1.82 | 35.32 | 0.006 | 8.95 | 1.44 | 55.79 | 0.019 |
| IgG2 < 10 mg/dl | 3.71 | 0.93 | 14.80 | 0.064 | 6.22 | 1.08 | 35.72 | 0.040 |
| ISC IgGAM = 0 | 13.79 | 2.61 | 72.98 | 0.002 | 17.53 | 2.23 | 137.57 | 0.006 |

#### III.2 Hospital mortality

when hospital mortality was analyzed, only ISC IgGAM showed differences for the proportion of survivors and non survivors between patients with low and high immunoglobulin levels (Table 4).

**Table 4:** Proportion of survivors and non survivors at the hospital in the group of patients with SOFA score < 8 depending on the immunoglobulin thresholds.

|  |  | *SOFA < 8: Hospital mortality* |  |  |  |  |  |
| --- | --- | --- | --- | --- | --- | --- | --- |
|  |  | *Patients with prior immunosuppression included (n=122)* |  |  | *Patients with prior immunosuppression excluded (n=99)* |  |  |
|  | Thresholds | Non survivors / Total "n" in each category | % | *p* | Non survivors / Total "n" in each category | % | *p* |
| **IgG (total)** | < 407 mg/dl | 10 / 31 | 32.3% | 0.119 | 7 / 24 | 29.2% | 0.209 |
|  | ≥ 407 mg/dl | 18 / 91 | 19.8% |  | 13 / 75 | 17.3% |  |
| **IgG1** | < 332 mg/dl | 10 / 30 | 33.3% | 0.119 | 7 / 24 | 29.2% | 0.209 |
|  | ≥ 332 mg/dl | 18 / 92 | 19.6% |  | 13 / 75 | 17.3% |  |
| **IgG2** | < 10 mg/dl | 11 / 39 | 28.2% | 0.344 | 9 / 32 | 28.1% | 0.175 |
|  | ≥ 10 mg/dl | 17 / 83 | 20.5% |  | 11 / 67 | 16.4% |  |
| **IgM** | < 43 mg/dl | 19 / 72 | 26.4% | 0.279 | 12 / 54 | 22.2% | 0.583 |
|  | ≥ 43 mg/dl | 9 / 50 | 18.0% |  | 8 / 45 | 17.8% |  |
| **IgA** | < 219 mg/dl | 14 / 49 | 28.6% | 0.226 | 10 / 39 | 25.6% | 0.277 |
|  | ≥ 219 mg/dl | 14 / 73 | 19.2% |  | 10 / 60 | 16.7% |  |
| **ISC IgGAM** | = 0 | 9 / 17 | 52.9% | 0.002 | 6 / 12 | 50.0% | 0.006 |
|  | = 1 | 19 / 105 | 18.1% |  | 14 / 87 | 16.1% |  |

**Table 5.** Multivariate logistic regression analysis to evaluate the association between immunoglobulins and the risk of mortality at the hospital in the group of patients with SOFA score < 8. Adjusting variables for *A.* were [age] [presence of prior immunosupression] [APACHE-II score] [microbiologically confirmed infection]. Adjusting variables for *B* were [APACHE-II score] [respiratory infection] [microbiologically confirmed infection].

| Immunoglobulins | SOFA < 8 |  |  |  |  |  |  |  |
| --- | --- | --- | --- | --- | --- | --- | --- | --- |
|  | A. Patients with prior immunosuppression included (n=122) |  |  |  | B. Patients with prior immunosuppression excluded (n=99) |  |  |  |
|  | OR | CI 95% |  | p | OR | CI 95% |  | p |
| ISC IgGAM = 0 | 7.98 | 1.94 | 32.87 | 0.004 | 5.63 | 1.13 | 28.11 | 0.035 |

Multivariate regression analysis for the ISCs combining two immunoglobulins were not performed since the vast majority of non survivors showing a “0” in the double ISCs showed also a “0” in the triple ISC (overlap with the triple score was 100% for ISC IgGM, of 90% for ISC IgGA and 82% for ISC IgMA).

### Potential influence of hemodilution on the results

interestingly, hemodilution had no effect on the results observed in the less severe group of patients, as evidenced by the absence of significant correlation in the Spearman Test between levels of immunoglobulins and albumin concentration in plasma (*p* > 0.05).

## Discussion

This study shows for the first time that the influence of endogenous immunoglobulin levels on the prognosis of patients with sepsis seems to be restricted to a subset of individuals presenting at the ICU with limited extent of organ failure. Our results evidenced also that the simultaneous presence of low levels of IgG, IgA and IgM was found to be a consistent predictor of both acute mortality (at the ICU) and post-acute mortality (at the hospital) in these patients, independently of the presence or absence of previous immunosupression. In contrast, the ability of total IgG or IgG1 to predict mortality was restricted to the acute period of the disease (ICU mortality). While the presence of patients with previous immunosupression did not alter the predictive ability of total IgG and IgG1, it modified that of IgG2, which was only able to predict ICU mortality in those patients with no previous immunosupression. These findings reinforce the superiority of the combined immunoglobulin score over immunoglobulins individually considered to identify sepsis patients at risk of poor outcomes [4].

In the era of precision medicine in sepsis [11] [12], defining the immunological state of the patient will be crucial to the success of any biological response modifier for sepsis [13] [14]. Our findings could contribute to personalize treatment with IVIG in this disease. Failure of IVIG in demonstrating clinical benefit in sepsis [15] could be explained by different factors concerning both the patient and the IVIG preparation, which have already been discussed elsewhere [1] [16]. Our results provide new clues to better design future trials with IVIG in sepsis and/or to improve data analysis from these trials. The more relevant would be that sepsis patients should not be considered as a homogenous population. The impact of IVIG of the outcome of the patients should be analyzed stratifying the patients by their levels of endogenous immunoglobulins & by the degree of disease severity at ICU admission. Quantification of immunoglobulin levels using nephelometry is a fast test which takes less than 2 hours. In turn, the SOFA score has widespread familiarity within the critical care community, making it useful to assess the degree of organ failure extent at ICU admission [7] [17]. In addition, our results provide a scientific rational to evaluate if IVIG preparations containing IgG, IgA, and IgM could be more effective for the treatment of sepsis than those containing exclusively IgG [18] [19] [20]. Interestingly, Kreymann KG *et al,* a metanalysis in 2007 of all randomized controlled studies published on polyvalent immunoglobulins for treatment of sepsis or septic shock, observed a strong protective trend in favor of an immunoglobulin preparation containing the three major immunoglobulin isotypes [21].

As a major limitation of our study, immunoglobulins were measured in samples already available from the GRECIA and the ABISS studies, and in consequence it was retrospective in nature. This makes that sample size was not calculated based on a predefined primary outcome, which may represent a source of bias. Another limitation was the absence of lactate registries in many patients, which precluded identifying those patients fulfilling the new definition of septic shock as proposed by the SEPSIS-3 consensus [8]. In consequence, new prospective studies should confirm the results obtained in this work.

## Conclusions

Results from this study suggest that endogenous immunoglobulin levels may have a different impact on the mortality risk of sepsis patients based on their severity. Future studies should be directed to investigate whether IVIG therapy may particularly benefit subsets of patients with moderate organ failure extent and low levels of the three immunoglobulin isotypes.

## List of abbreviations

ABISS-Edusepsis: AntiBiotic Intervention in Severe Sepsis
APACHE-II: Acute Physiology and Chronic Health Evaluation II score
GRECIA: Grupo de Estudios y Análisis en Cuidados Intensivos.
IVIG: intravenous immunoglobulin
ISC IgGAM: immunoscore combining IgG, IgA, IgM
ISC IgGM: immunoscore combining IgG, IgM
ISC IgGA: immunoscore combining IgG, IgA
ISC IgMA: immunoscore combining IgM, IgA.
SBITs: Score-based immunoglobulin G therapy of patients with sepsis
SOFA: Sequential Organ Failure Assessment score

## Ethics approval and consent to participate

Written informed consent was obtained directly from all patients, or their legal representative, before enrolment. Scientific and ethical approval of the study protocol was obtained from the ethical committees for clinical research of all participating hospitals. The study has been performed in accordance with the ethical standards laid down in the 1964 Declaration of Helsinki and its later amendments.

## Consent for publication

Not applicable

## Availability of data and materials

The datasets used during the current study are available from the corresponding author on reasonable request.

## Competing interests

The authors declare that they have no competing interests

## Funding

The GRECIA study was supported by “Proyectos de Investigación en Biomedicina, Consejería de Sanidad, JCYL (grant code BOCYL-D-26072010)”. The authors thank also Instituto de Salud Carlos III for their financial support (grant code EMER 07/050, ISCIII-FIS-PI12-01815). The funding agencies were not involved neither in the design of the study and collection, not in the analysis, interpretation of data or manuscript writing

## Authors’ contributions

IML, JFBM designed the study and wrote the article; AMB, RF, AA, JSV, LL, DAO, APB, RHM, BS, PM, AML, PGO, EA, ET, JB, GRECIA group recruited the patients and samples and critically reviewed the manuscript; ARF and WT drafted the article tables and figures, and critically reviewed the manuscript

## Acknowledgements

We want to thank the role of Gemma Goma as research nurse in charge of sample and data collection for the ABISS-Edusepsis study.

Researchers of the GRECIA group (Coordinator: Jesús Blanco)
Marta María García-García (Hospital Universitario Río Hortega, Valladolid, Spain); M^a^ Jesús López Pueyo, Jose Antonio Fernandez Ratero, Miguel Martinez Barrios, Fernando Callejo Torre, Sergio Ossa Echeverri (Hospital General Yagüe, Burgos, Spain); Demetrio Carriedo Ule, Ana Ma Domínguez Berrot, Fco Javier Díaz Domínguez (Complejo Hospitalario de León, Spain); Susana Moradillo (Hospital Río Carrión, Palencia, Spain); Braulio Alvarez Martínez (Hospital del Bierzo, Ponferrada, Spain); Noelia Albalá, Juan Carlos Ballesteros, Marta Paz Perez, Elena Perez Losada (Hospital Clínico Universitario de Salamanca, Spain); Santiago Macías, Rafael Pajares García, Noelia Recio García Cervigón (Hospital General de Segovia Spain); M^a^ Mar Gobernado Serrano, M^a^ José Fernández Calavia, Daniel Moreno Torres (Complejo Hospitalario de Soria, Spain); Concha Tarancón, Teresa Loreto, Priscila Carcelen (Hospital Virgen de la Concha, Spain), Rafael Cítores y Francisco Gandía (Hospital Clínico Universitario de Valladolid).

## Additional files

**Additional file 1: Mean survival time (days) in the SOFA < 8 group based on immunoglobulin thresholds**. Differences between groups were assessed using the log-rank test. Δ (days) represents [(mean survival time in patients with levels of immunoglobulin above the threshold) – (mean survival time in patients with levels of immunoglobulin below the threshold)]. Time was censored at 28 days following ICU admission.

## References

1. Almansa R, Tamayo E, Andaluz-Ojeda D, Nogales L, Blanco J, Eiros JM, et al. The original sins of clinical trials with intravenous immunoglobulins in sepsis. Crit. Care Lond. Engl. 2015;19:90.

2. Shankar-Hari M, Culshaw N, Post B, Tamayo E, Andaluz-Ojeda D, Bermejo-Martín JF, et al. Endogenous IgG hypogammaglobulinaemia in critically ill adults with sepsis: systematic review and meta-analysis. Intensive Care Med. 2015;41:1393–401.

3. Dietz S, Lautenschläger C, Müller-Werdan U, Pilz G, Fraunberger P, Päsler M, et al. Serum IgG levels and mortality in patients with severe sepsis and septic shock: The SBITS data. Med. Klin. Intensivmed. Notfallmedizin. 2016;

4. Bermejo-Martín JF, Rodriguez-Fernandez A, Herrán-Monge R, Andaluz-Ojeda D, Muriel-Bombín A, Merino P, et al. Immunoglobulins IgG1, IgM and IgA: a synergistic team influencing survival in sepsis. J. Intern. Med. 2014;276:404–12.

5. Bermejo Martin, Jesus F, Giamarellos-Bourboulis, Evangelos J. Endogenous immunoglobulins and sepsis: New perspectives for guiding replacement therapies. Int. J. Antimicrob. Agents. 2015;

6. Andaluz-Ojeda D, Nguyen HB, Meunier-Beillard N, Cicuéndez R, Quenot J-P, Calvo D, et al. Superior accuracy of mid-regional proadrenomedullin for mortality prediction in sepsis with varying levels of illness severity. Ann. Intensive Care. 2017;7:15.

7. Singer M, Deutschman CS, Seymour CW, Shankar-Hari M, Annane D, Bauer M, et al. The Third International Consensus Definitions for Sepsis and Septic Shock (Sepsis-3). JAMA. 2016;315:801–10.

8. Shankar-Hari M, Phillips GS, Levy ML, Seymour CW, Liu VX, Deutschman CS, et al. Developing a New Definition and Assessing New Clinical Criteria for Septic Shock: For the Third International Consensus Definitions for Sepsis and Septic Shock (Sepsis-3). JAMA. 2016;315:775–87.

9. Herrán-Monge R, Muriel-Bombín A, García-García MM, Merino-García PA, Cítores-González R, Fernández-Ratero JA, et al. Mortality Reduction and Long-Term Compliance with Surviving Sepsis Campaign: A Nationwide Multicenter Study. Shock Augusta Ga. 2016;45:598–606.

10. Sánchez B, Ferrer R, Suarez D, Romay E, Piacentini E, Gomà G, et al. Declining mortality due to severe sepsis and septic shock in Spanish intensive care units: A two-cohort study in 2005 and 2011. Med. Intensiva. 2017;41:28–37.

11. Bermejo-Martin JF, Andaluz-Ojeda D, Almansa R, Gandía F, Gómez-Herreras JI, Gomez-Sanchez E, et al. Defining immunological dysfunction in sepsis: A requisite tool for precision medicine. J. Infect. 2016;72:525–36.

12. Bermejo-Martin JF, Tamayo E, Andaluz-Ojeda D, Martín-Fernández M, Almansa R. Characterizing Systemic Immune Dysfunction Syndrome to Fill in the Gaps of SEPSIS-2 and SEPSIS-3 Definitions. Chest. 2017;151:518–9.

13. Almansa R, Wain J, Tamayo E, Andaluz-Ojeda D, Martin-Loeches I, Ramirez P, et al. Immunological monitoring to prevent and treat sepsis. Crit. Care Lond. Engl. 2013;17:109.

14. Hotchkiss RS, Moldawer LL, Opal SM, Reinhart K, Turnbull IR, Vincent J-L. Sepsis and septic shock. Nat. Rev. Dis. Primer. 2016;2:16045.

15. Alejandria MM, Lansang MAD, Dans LF, Mantaring JB. Intravenous immunoglobulin for treating sepsis, severe sepsis and septic shock. Cochrane Database Syst. Rev. 2013;CD001090.

16. Shankar-Hari M, Spencer J, Sewell WA, Rowan KM, Singer M. Bench-to-bedside review: Immunoglobulin therapy for sepsis - biological plausibility from a critical care perspective. Crit. Care Lond. Engl. 2012;16:206.

17. Vincent JL, de Mendonça A, Cantraine F, Moreno R, Takala J, Suter PM, et al. Use of the SOFA score to assess the incidence of organ dysfunction/failure in intensive care units: results of a multicenter, prospective study. Working group on “sepsis-related problems” of the European Society of Intensive Care Medicine. Crit. Care Med. 1998;26:1793–800.

18. Neilson AR, Burchardi H, Schneider H. Cost-effectiveness of immunoglobulin M-enriched immunoglobulin (Pentaglobin) in the treatment of severe sepsis and septic shock. J. Crit. Care. 2005;20:239–49.

19. Welte T, Dellinger RP, Ebelt H, Ferrer M, Opal SM, Schliephake DE, et al. Concept for a study design in patients with severe community-acquired pneumonia: A randomised controlled trial with a novel IGM-enriched immunoglobulin preparation - The CIGMA study. Respir. Med. 2015;109:758–67.

20. Giamarellos-Bourboulis EJ, Tziolos N, Routsi C, Katsenos C, Tsangaris I, Pneumatikos I, et al. Improving outcomes of severe infections by multidrug-resistant pathogens with polyclonal IgM-enriched immunoglobulins. Clin. Microbiol. Infect. Off. Publ. Eur. Soc. Clin. Microbiol. Infect. Dis. 2016;22:499–506.

21. Kreymann KG, de Heer G, Nierhaus A, Kluge S. Use of polyclonal immunoglobulins as adjunctive therapy for sepsis or septic shock. Crit. Care Med. 2007;35:2677–85.

